# Trafficking and translation of mRNA in osteocyte dendrites

**DOI:** 10.1101/2024.10.01.616162

**Authors:** Courtney M. Mazur, Parthena E. Kotsalidis, Christian D. Castro Andrade, Mingrui Wu, Xiao Wang, J. Matthew Taliaferro, Marc N. Wein

## Abstract

Osteocyte dendrites extend great distances through canaliculi in bone supported by an extensive cytoskeletal network. The formation and maintenance of osteocyte dendrites is closely linked to bone health, yet the mechanisms regulating these processes are poorly understood. Here we tested the hypothesis that osteocytes transport mRNA to their dendrites for local translation as a mechanism of regulating the formation and maintenance of these subcellular structures. Using molecular and imaging approaches, we demonstrated that a subset of mRNAs are enriched in Ocy454 osteocyte-like cell dendrites. To understand how certain mRNAs are directed to dendrites, we performed a massively parallel reporter assay with 7115 oligonucleotide sequences designed from the 3’ untranslated regions of eight dendrite-enriched transcripts. We identified localization sequences within some 3’UTRs that were both sufficient and necessary for trafficking to osteocyte dendrites, indicating that localization directions are specifically encoded in enriched transcripts. Furthermore, we found that dendrites contain the elements necessary for local translation, and trafficked transcripts were engaged in active translation within dendrites. Together this work describes a molecular regulatory mechanism novel to osteocyte biology with many potential consequences for the spatiotemporal control of the cytoskeleton and dendrites.

## INTRODUCTION

Osteocytes are bone embedded cells that extend long dendritic projections in all directions throughout mineralized bone. Dendrites develop as osteoblasts terminally differentiate and embed in newly deposited bone, and abundance and length of dendrites are associated with osteocyte health (Moharrer and Boerckel 2021; J. S. Wang and Wein 2022). Healthy osteocytes are essential for organized bone matrix and strong bone through their functions in mechanosensation, perilacunar/canalicular remodeling, and production of signaling molecules that govern osteoblast and osteoclast activity on bone surfaces (Dallas, Prideaux, and Bonewald 2013). Defects in the osteocyte dendrite network are frequently observed during aging (Tiede-Lewis et al. 2017) and in genetic or acquired skeletal diseases (J. S. Wang et al. 2021; Fowler et al. 2017). The formation and maintenance of osteocyte dendrites is of great importance for overall bone health, yet little is known about the molecular mechanisms used by osteocytes to regulate their dendrites during skeletal health and disease.

Osteocyte dendrites stretch tens of microns through bone, supported by a cytoskeleton of actin filaments and microtubules. Loss of many individual factors (Sp7, Tgfbr2, Mmp13, Pdpn, and others (J. S. Wang et al. 2021; Dole et al. 2017; Mazur et al. 2019; Staines et al. 2017)) results in osteocyte dendrite defects due to cells being unable to support dendrite formation (cytoskeletal defects) and/or difficulty in creating space in the bone for dendrites to fill (protease or bone mineral defects). We focused on the processes that govern biogenesis of the osteocyte dendrite cytoskeleton.

In other cell types with specialized subcellular regions, the local enrichment of select mRNAs contributes to their morphology and function. Proactive transport of specific subsets of mRNAs allows for local translation, where one transcript can serve as template for multiple copies of a protein that are immediately localized at their site of function (Zappulo et al. 2017; Engel et al. 2020; Biever et al. 2020). Thus, osteocyte dendrites might be maintained by local protein synthesis rather than transporting proteins needed for the dendrite cytoskeleton across many microns. We therefore hypothesized that osteocytes transport mRNA to their dendrites for local translation as a molecular regulatory mechanism supporting their morphology and function.

Here we show that Ocy454 cell osteocyte dendrites contain enrichment of a specific subset of transcripts. Localization sequences encoded in the 3’UTRs are necessary and sufficient for the transport of many of these mRNAs to dendrites. Dendrites also contain the elements necessary for local translation, and all the dendrite-enriched transcripts tested were engaged in active translation in association with ribosomes distant from the nucleus. Together, this work introduces a molecular regulatory mechanism novel to osteocyte biology and provides a new source of skeletal disease targets through mislocalization.

## RESULTS

### Identification of osteocyte dendrite-enriched mRNAs

We adapted subcellular fractionation methods to identify and quantify transcripts present in osteocyte dendrites (Arora et al. 2021). Osteocyte-like Ocy454 cells bearing a temperature sensitive large T antigen (active at 33ºC and inactive at 37ºC (Spatz et al. 2015; Wein et al. 2015)) were grown on microporous membranes, which support the cell body while allowing projections to grow through and below 1 µm pores (Figure 1A, B). At confluence (Day 0), cells are switched to 37ºC, which slows their growth and begins the differentiation program. Between 0 and 7 days of differentiation, the number of phalloidin-stained dendrites extending through the membrane increases (Supp Figure 1A). To fractionate, we gently scrape the soma from the membranes and collect the cell suspension, followed by direct lysis of the dendrites from the membrane for RNA or protein isolation (Figure 1A). Imaging of membranes after scraping confirms that DAPI-stained nuclei are depleted and phalloidin-stained cell projections are retained within the membrane (Figure 1B). Dendrite protein lysates contain b-actin but not Histone H3, indicating removal of nuclei from the dendrite fraction (Supp Figure 1B).

**Figure 1:**
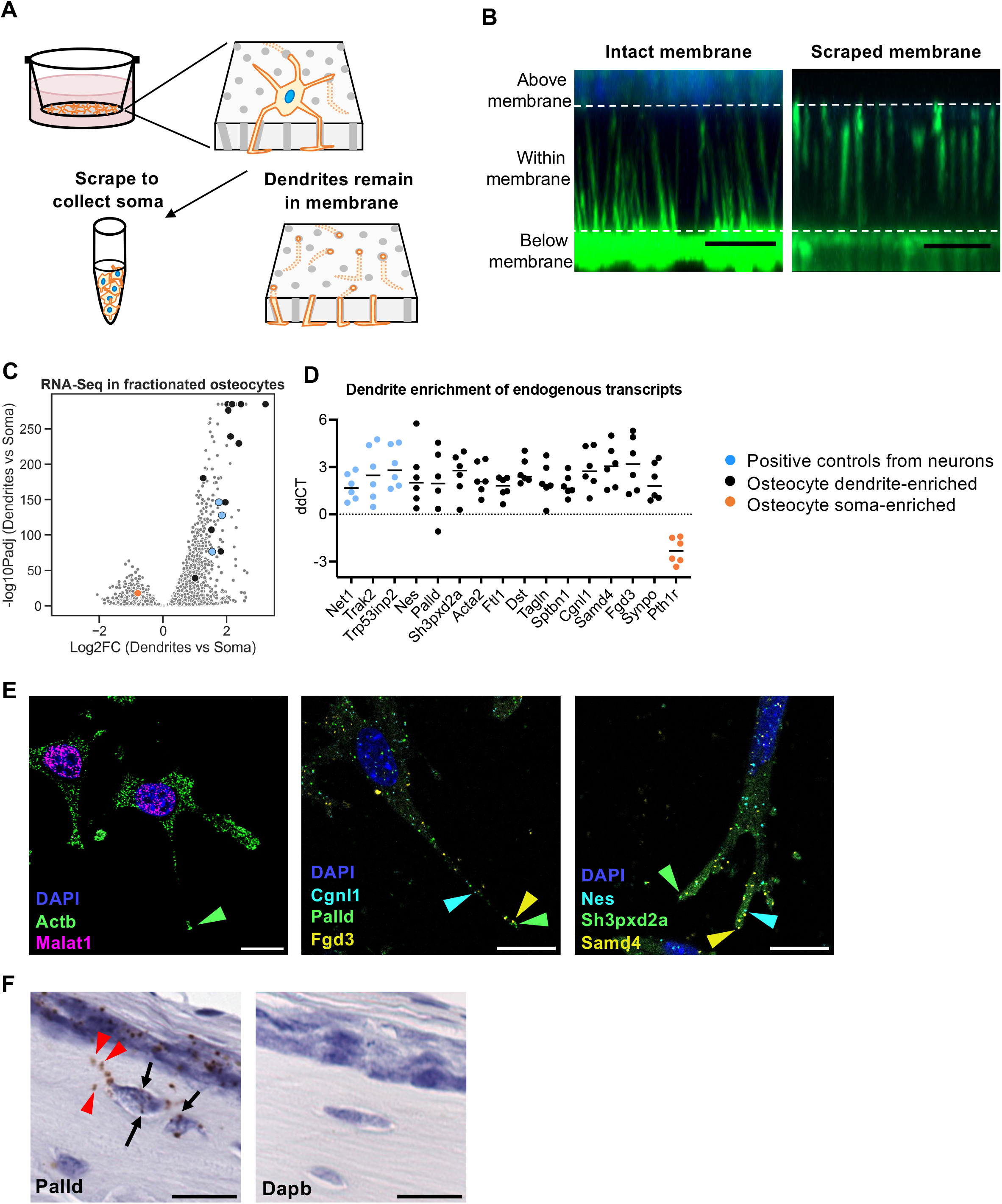
Osteocyte dendrites contain selected mRNAs. A) Schematic illustrating mechanical fractionation of osteocytes using microporous membranes. B) Side view of Z-stack images of F-actin stained dendrites extending through microporous membranes. Dashed lines indicate the top and bottom of the membrane. Scale bars = 10 μm. C) Volcano plot showing dendrite/soma enrichment for all expressed genes in Ocy454 cells based on RNA-Seq, n=3 biologic replicates. D) qRT-PCR for endogenous transcript enrichment in dendrites compared to soma, calculated relative to *Pgk1* housekeeping gene, n=6. E) STARmap for indicated endogenous mRNAs. Arrows indicate location of example puncta in dendrites. Scale bars = 20 μm. F) RNAscope in 4-week old mouse tibia. Scale bars = 20 μm.

We collected RNA from soma and dendrite fractions and performed RNA sequencing. This revealed that dendrites contain thousands of mRNA transcripts, with over 400 mRNAs significantly enriched in dendrites versus cell bodies (log_2_FC≥1, p_adj_<0.01, Figure 1C, Supp Figure 1D). Gene ontology analysis indicates dendrite enrichment of transcripts encoding ribosomal proteins and electron transport chain components, consistent with previous findings in neurons (Goering et al. 2020), as well as cytoskeletal proteins (Supp Figure 1E). Transcripts previously reported as enriched in neuronal dendrites – *Net1, Trak2*, and *Trp53inp2* –also showed enrichment in osteocyte dendrites (Arora et al. 2022).

We identified *Hprt* and *Pgk1* as potential housekeeping genes without enrichment in either fraction and performed RT-qPCR with equal RNA/cDNA input from independent samples. We found that *Pgk1* levels were the most consistently expressed across fractions. We then selected dendrite-enriched genes from the RNA-seq for confirmation by RT-qPCR using *Pgk1* as a housekeeping gene. We selected *Ftl1, Nes, Palld, Sh3pxd2a, Acta2, Dst*, and *Tagln* based on the great magnitude of their enrichment by RNA-seq. *Cgnl1* and *Synpo* were selected because they are not typically expressed in neurons and did not appear in prior studies of mRNA localization (Middleton, Eberwine, and Kim 2019). *Fgd3, Samd4*, and *Sptbn1* are expressed in neurons but show more dendrite enrichment in our osteocytes than in published data from fractionated CAD neurons (Taliaferro et al. 2016). Using RT-qPCR, these genes were all confirmed as dendrite-enriched in independent samples (Figure 1D).

In contrast, transcripts enriched in the cell body fraction included *Runx2* and *Sp7* (transcription factors), *Dmp1* and *Ihh* (secreted proteins), and *Pth1r* and *Bambi* (transmembrane proteins). These findings are consistent with the requirement for mRNAs encoding trans-membrane or secreted proteins to be translated in peri-nuclear rough endoplasmic reticulum which remains in the soma fraction in our workflow (Supp Figure 1C).

Next, we used STARmap (X. Wang et al. 2018) to visualize mRNA localization in 2D cultured Ocy454 cell dendrites. As expected, abundant *Actb* puncta were located throughout cell projections, whereas the non-coding RNA *Malat1* was confined to the nucleus (Figure 1E), validating the use of STARmap in Ocy454 cells. We then probed the location of osteocyte dendrite-enriched mRNAs (*Cgnl1, Palld, Fgd3, Nes, Sh3pxd2a*, and *Samd4*), and found puncta throughout dendrites as far as 60 μm from the nucleus (Figure 1E). Critically, we also noted *Palld* mRNA within dendritic structures of murine bone-embedded osteocytes *in vivo* (Figure 1F).

Together these results demonstrate the presence of specific mRNAs within osteocyte dendritic projections.

### Translation in osteocyte dendrites

Having identified distinct mRNAs within osteocyte dendrites, we next asked if translational machinery components are present in these structures. Based on our RNA-seq results, mRNAs for genes encoding 60S and 40S ribosomal subunit proteins were enriched in dendrites (Figure 2A). Rpl26 protein is present in soma and dendrites of fractionated Ocy454 cells (Supp Figure 1B). We also detected 18s, 28s and 5.8s ribosomal RNAs in dendrite RNA fractions by RT-qPCR with significant enrichment versus soma (Figure 2B). Furthermore, 18s rRNA was visible throughout osteocyte dendrites within mouse bone (Figure 2C). Therefore, osteocyte dendrites contain components necessary for local translation of mRNA to protein.

**Figure 2:**
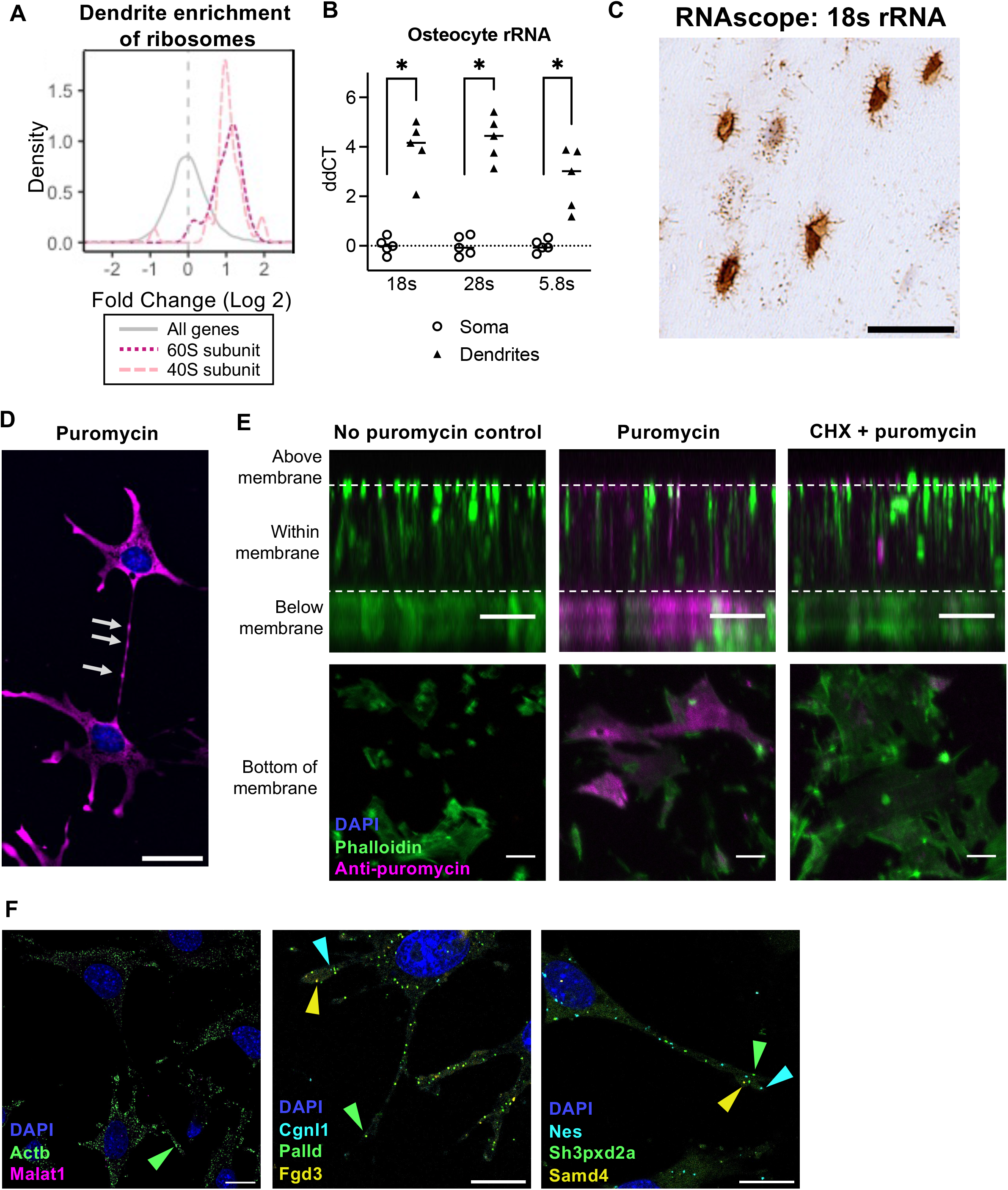
Osteocyte dendrites translate mRNA into protein. A) Dendrite enrichment values from RNA-Seq for all genes (grey), and genes encoding large and small ribosomal subunit proteins (pink). B) qRT-PCR for rRNA in dendrites and soma calculated relative to *Pgk1* housekeeping gene, n=5, *p<0.05. C) RNAscope in 4-week old mouse tibia. Scale bar = 20 μm. D) Anti-puromycin immunofluorescence (magenta) after 10 minutes puromycin treatment in D) Ocy454 cells grown in 2D. Arrows indicate puromycylated peptides in dendrites. Scale bar = 20 μm. E) Side view of Z-stack images of dendrites extending through microporous membranes (top) and bottom view of membranes as dendrites emerge from pores (bottom) at day 2 of differentiation, stained for F-actin (green) and anti-puromycin immunofluorescence (magenta) after 30 minutes of the indicated treatments. Dashed lines indicate the top and bottom of the membrane. Scale bars = 10 μm. F) RIBOmap for indicated mRNAs. Arrows indicate location of example puncta in dendrites. Scale bars = 20 μm.

To visualize nascent polypeptides, we pulsed Ocy454 cells with puromycin for 5-10 minutes and then performed immunofluorescence to detect puromycylated proteins (Schmidt et al. 2009). Within this short time frame, newly synthesized puromycylated proteins were visible within osteocyte dendrites more than 20 microns from the nucleus in a cycloheximide-sensitive manner (Figure 2D, Supp Figure 2A). Since, in theory, proteins could be synthesized in the soma and then transported this distance during the experimental period, we grew Ocy454 cells on transwell membranes and scraped away cell bodies just prior to adding puromycin (Torre and Steward 1992). Again, nascent puromycylated polypeptides were detected within isolated osteocyte dendrites using both immunostaining (Figure 2E) and immunoblotting (Supp Figure 2B, C), confirming that translation in these structures can occur independent of cell bodies.

To determine whether osteocyte dendrite-localized mRNAs of interest are translated into protein in osteocyte dendrites, we used RIBOmap spatial *in situ* translational profiling (Zeng et al. 2023). Probes targeting specific mRNAs and 18s rRNA must hybridize in close proximity to initiate rolling circle amplification, and thus only ribosome-associated mRNA is detected as a fluorescent puncta. In 2D cultured Ocy454 cells, we observed RIBOmap *Actb* puncta throughout the cell and extending to the ends of dendrites, demonstrating that *Actb* mRNAs in osteocyte dendrites are engaged with ribosomes and undergoing active translation (Figure 2F). Almost no puncta were visible for the non-coding RNA *Malat1*. RIBOmap puncta indicating the location of actively-translating *Cgnl1, Palld, Fgd3, Nes, Sh3pxd2a*, and *Samd4* mRNA were also distributed throughout dendrites and over 50 μm from the nucleus. This demonstrates that specific dendrite-enriched mRNA transcripts engaged in translation in dendrites.

### Some mRNAs are trafficked to osteocyte dendrites through localization sequences within their 3’ UTR

Next, we sought to understand how a subset of the osteocyte transcriptome achieves dendrite localization. Multiple mechanisms have been proposed to explain how mRNAs within many organisms and cell types become subcellularly enriched, including sequestration of RNA, selective protection or degradation, or formation of a complex with motor proteins for active transport (Chekulaeva 2024). One prominent mechanism involves RNA binding proteins that recognize 3’ untranslated region (3’UTR) ‘localization sequences’ to mediate the interaction with motor proteins. Thus, we tested the hypothesis that osteocyte dendrite-enriched transcripts contain localization sequences in their 3’UTRs.

We used a heterologous reporter system in which the 3’UTR of a dendrite-enriched mRNA is cloned downstream of the Firefly luciferase (FLuc) coding sequence (Arora et al. 2022). The Renilla luciferase (RLuc) coding sequence without an inserted UTR sequence serves as a control expressed from the same doxycycline-inducible promoter. Ocy454 cells were modified to contain asymmetric LoxP acceptor sites in the Rosa26 locus, and each expression construct was stably integrated by Cre/LoxP-mediated recombination (Khandelia, Yap, and Makeyev 2011). Cell lines expressing a single 3’UTR reporter were each grown on microporous membranes, treated with doxycycline, and subjected to soma/dendrite fractionation and RNA isolation. FLuc and RLuc expression were then measured by RT-qPCR in dendrites and cell bodies to determine whether the presence of each candidate 3’UTR is sufficient to confer dendrite enrichment of the FLuc coding sequence.

For 8 of 12 dendrite-enriched transcript 3’UTRs that were tested, the 3’UTR sequence was sufficient to confer dendrite FLuc enrichment (Figure 3A). Therefore, these eight 3’UTRs contain a localization sequence. As positive controls, we used the 3’UTRs of *Net1, Trak2*, and *Trp53inp2*, which contain active neurite-targeting localization sequences in neurons (Arora et al. 2022). As a negative control, we used the 3’UTR of cell body-enriched *Pth1r* mRNA, which did not promote dendrite localization. For dendrite-enriched mRNAs with 3’UTRs that did not promote FLuc localization, it is possible that localization information lies within the 5’UTR, coding sequence, or alternative splice forms of the 3’UTR. These results confirm regulation of RNA localization via 3’UTRs for some of our genes of interest, however they do not identify the specific localization sequence within each entire 3’UTR.

**Figure 3:**
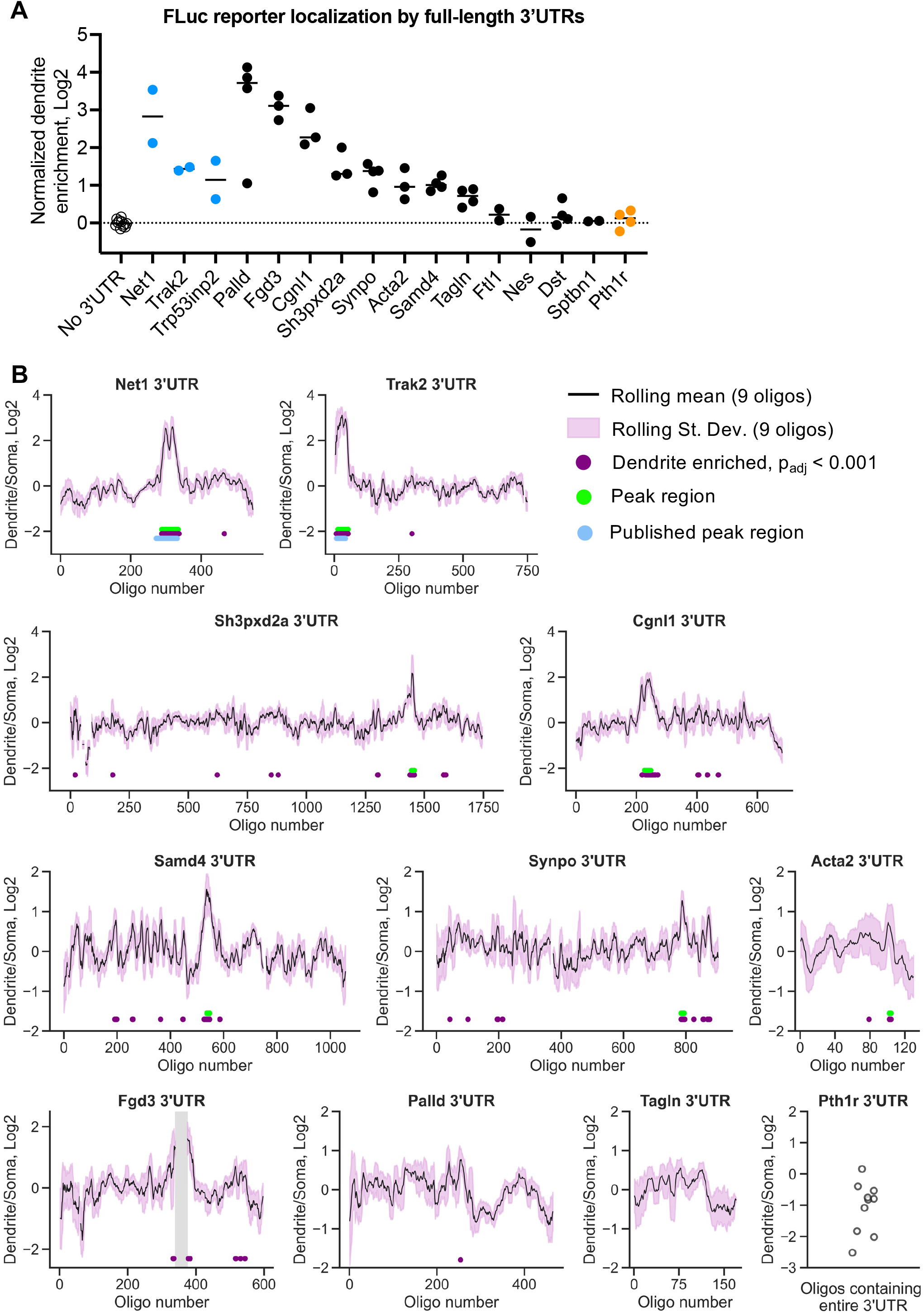
Identification of 3’UTR localization sequences responsible for dendrite enrichment of selected transcripts. A) Dendrite enrichment of FLuc fused to the 3’UTR of the indicated genes relative to RLuc. The localization of FLuc without an added 3’UTR was used as a normalization control in each experiment. B) Dendrite enrichments of oligos fused to the GFP reporter in the MPRA in order of their location within each 3’UTR. Lines represent a centered sliding average of nine oligos, and the ribbon represents the standard deviation for the sliding average. Purple dots below the lines indicate significant dendrite enrichment (Log2FC>0, p_adj_<0.001), and green dots indicate peaks of dendrite localization activity. In the *Fgd3* 3’UTR, the grey bar indicates the 39 consecutive oligos below the detection threshold for analysis. The entire *Pth1r* 3’UTR (169 nt) was represented in ten oligos each indicated with a single point.

### Precise mapping of localization sequences via massively parallel reporter assay

To precisely identify localization sequences within 3’UTRs sufficient to confer dendrite localization, we designed a massively parallel reporter assay (MPRA) covering the 3’UTRs of *Palld, Fgd3, Sh3pxd2a, Cgnl1, Samd4, Synpo, Acta2*, and *Tagln*, plus the 3’UTRs of *Net1, Trak2*, and *Pth1r* as controls. Each 3’UTR sequence was broken into 260 nucleotide (nt) fragments tiled at 4 nt intervals. The resulting library of 7115 oligos was cloned into two expression sites in a reporter plasmid: once following the FLuc coding sequence, and once following the GFP coding sequence, both under control of doxycycline-inducible promoters. Use of two independent sites affords the opportunity for a synchronous replicate within this initial MPRA. To ensure representation of all 7115 oligos in the pooled plasmid library, we conducted NGS and aligned reads to the oligonucleotide library. This revealed that our cloned plasmid pool contained more than 99.8% of designed oligos at both cloning sites (Supp Figure 3A).

As with the full length 3’UTR reporter constructs, we aimed to integrate the pooled plasmid library into the Rosa26 locus of Ocy454 cells so that each cell would stably express one plasmid from the library. Final integration efficiency of the library depends on both transfection efficiency of the plasmid into Ocy454 cells and Cre/LoxP recombination efficiency. To determine net integration efficiency in our cells, we transfected a plasmid library containing 15 nt randomers along with a Cre-expressing plasmid into LoxP-modified Ocy454 cells and performed targeted RNA-seq to measure the number of unique integrated randomers. When 6*10^5^ cells were transfected, we detected 9*10^4^ unique integration events (15% integration efficiency). Therefore, we performed transfections for the MPRA using 5*10^6^ cells to obtain at least 100 independent integrations of all 7115 oligos.

We collected RNA from the resulting pool of MPRA Ocy454 cells after doxycycline treatment to verify reporter integration. We generated cDNA libraries using reverse transcription primers that targeted the FLuc and GFP reporter sites and incorporated a unique molecular identifier (UMI) for each reverse transcription event. Using this targeted RNA-Seq approach, we detected at least 10 UMIs for 99.4% of oligos at both cloning sites (Supp Figure 3A).

We then grew MPRA Ocy454 cells on transwell membranes, treated with doxycycline, performed soma/dendrite fractionation, and used the same targeted RNA-Seq approach to measure the relative abundance of each oligo in dendrite and soma fractions. Oligo abundance within the samples clustered by compartment for three out of four biologic replicates. We therefore continued analysis with only the three paired samples with highly similar oligo abundance within compartments. We calculated relative oligo enrichment in dendrites for each oligo at each reporter site using DESeq2. With an adjusted p-value cutoff of less than 0.01, we found 369 oligos with significant enrichment in dendrites and 368 oligos that were enriched in soma using the GFP reporter. Oligos that were dendrite-enriched in the GFP reporter context also had significantly higher dendrite enrichment when analyzed in the FLuc reporter context (Supp Figure 3B). Therefore, oligo localization activity is independent of the reporter context. Dendrite-enriched oligos had significantly higher A/G content than soma or non-localized oligos (Supp Figure 3C), in line with prior studies of 3’UTR-driven localization (Arora et al. 2022).

We next sorted oligo enrichment results from the GFP reporter by their position within each 3’UTR (Figure 3B). With a more stringent cutoff of p_adj_<0.001, peaks of dendrite localization activity are clearly visible within most 3’UTRs. The 3’UTRs of *Trak2* and *Net1* had previously been investigated in CAD neurons using this approach and showed nearly equivalent peaks in osteocytes, suggesting conservation of this regulatory mechanism across cell types for these two transcripts (Arora et al. 2022). The 3’UTRs of *Sh3pxd2a, Cgnl1, Samd4, Synpo*, and *Acta2* had peaks of 3-22 adjacent oligos that caused significant dendrite localization (p_adj_<0.001). Since oligos in each peak overlap, multiple independent tests contribute to the conclusion that a localization sequence is present in the peak region. The A/G content within each peak region is higher than in the surrounding sequence of each 3’UTR (Supp Figure 3D). The 3’UTR of soma-enriched *Pth1r* mRNA (169 nt) was contained within ten oligos, and as expected, none were significantly dendrite-enriched.

While the full-length 3’UTRs of *Palld* (1919 nt) and *Tagln* (752 nt) confer dendrite localization to a heterologous reporter, no 260 nt oligos within these 3’UTRs were dendrite-enriched. In these cases, the localization sequence may distributed over more than 260 nt or depend on the secondary structure provided by the greater sequence context for activity. In the 3’UTR of *Fgd3*, 39 adjacent oligos were not sufficiently detected in the final analysis to calculate dendrite enrichment. This is likely due to high GC content of oligos in this region (>60%) affecting their amplification during library preparation for targeted RNA-seq. However, oligos on both sides flanking the missing region were significantly dendrite-enriched, suggesting that the missing *Fgd3* 3’UTR region may contain additional dendrite-targeting oligos that could be studied outside of the MPRA.

### Validation of localization sequences identified by MPRA

To verify the function of the peak regions identified in the MPRA, we created individual reporter plasmids bearing a single 260 nt dendrite-enriched oligo from the peak regions of *Sh3pxd2a, Cgnl1, Samd4, Synpo*, and *Acta2* cloned downstream of the FLuc coding sequence. For *Fgd3*, we chose a single dendrite-enriched oligo preceding the GC-rich region absent from the MPRA analysis, a single oligo centered within the missing region, and the 412 nt union of the 39 oligos absent from the MPRA analysis.

Five of the individual oligo sequences and the *Fgd3* missing oligo union were sufficient to promote significant FLuc dendrite enrichment (Figure 4A). For *Sh3pxd2a* and *Synpo*, the activity of one peak oligo was comparable to that of the full-length 3’UTR. For *Cgnl1, Samd4*, and two of the sequences from *Fgd3*, dendrite localization occurred albeit to a lesser extent than what was observed with the gene’s full-length 3’UTR. We also created reporter plasmids with 500-668 nt fragments of the *Palld* 3’UTR, none of which caused dendrite enrichment to the same extent as the full-length 3’UTR (Supp Figure 4A). Together, this shows strong regulatory activity of the peak sequences identified by MPRA and suggests that, for some transcripts, the full 3’UTR can contain multiple elements that work together to modify the degree of subcellular localization.

**Figure 4:**
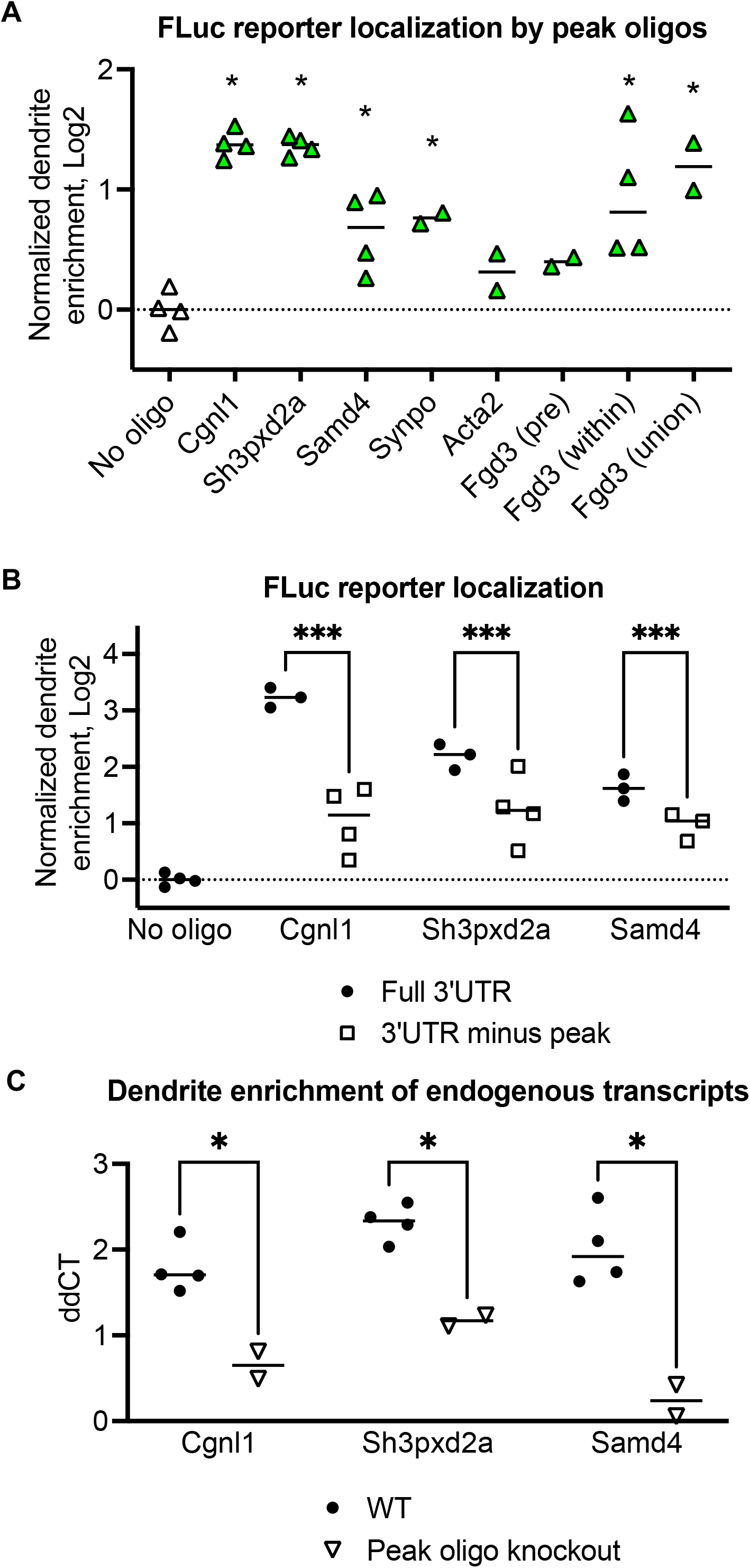
Localization sequences identified by MPRA are necessary and sufficient for dendrite enrichment. A) Dendrite enrichment of FLuc fused to one peak oligo (260 nt) identified from the MPRA relative to RLuc. For Fgd3, two individual oligos were tested, as well as the 412 nt union of the 39 consecutive oligos absent from the MPRA analysis. The localization of FLuc without any added localization sequence was used as a normalization control in each experiment. *p<0.05 compared to no oligo control. B) Dendrite enrichment of FLuc fused to full 3’UTRs or 3’UTRs lacking the sequences of the peak 260 nt oligo relative to RLuc. The localization of FLuc without any added localization sequence was used as a normalization control in each experiment. ***p<0.001. C) qRT-PCR for endogenous transcript enrichment in dendrites compared to soma, calculated relative to *Pgk1* housekeeping gene, for wild-type cells and cells with deletion of the localization sequence from the genome. *p<0.05.

Having demonstrated that MPRA-identified localization sequences are sufficient to confer dendrite targeting in the heterologous reporter system, we next asked if these sequences are necessary for dendrite mRNA localization. First, we generated reporter constructs with the 3’UTRs of *Sh3pxd2a, Cgnl1*, and *Samd4* lacking their respective peak 260 nt oligo sequences. For all three sequences, removal of the peak oligo reduced reporter localization relative to the full 3’UTR (Figure 4B). Next, the corresponding regions of these endogenous transcripts were deleted by CRISPR/Cas9 using paired sgRNAs flanking the newly-identified localization sequences. Single cell clones with confirmed bi-allelic deletion (Supp Figure 4B) were grown on transwell membranes and endogenous transcript dendrite localization was assessed by RT-qPCR. Deletion of the peak region from the genome also reduced dendrite enrichment of the corresponding endogenous mRNAs (Figure 4C). Together, this indicates that localization sequences identified from the MPRA are necessary for full localization activity of the corresponding 3’UTRs and transcripts.

## DISCUSSION

Here we demonstrated that osteocytes employ directed, specific mRNA trafficking to their dendrites. Using sequencing- and imaging-based subcellular transcriptomic strategies, we showed that a subset of mRNAs are enriched in osteocyte dendrites under control of localization sequences within their 3’UTRs. In a heterologous reporter system, 3’UTR localization sequences were both necessary and sufficient for trafficking to osteocyte dendrites. Puromycylation and RIBOmap approaches support the model that new protein translation occurs in osteocyte dendrites, including translation from dendrite-enriched mRNAs. These data illuminate mRNA trafficking as a novel molecular regulatory mechanism in osteocytes. Given the central role for these cells in skeletal biology and the key roles of dendrites in osteocyte function, our findings highlight mRNA trafficking as a novel mechanism with enormous potential importance in bone homeostasis.

Studies of osteocyte biology typically focus on single genes or proteins that impact cell shape or function. Instead, we decided to investigate the molecular regulatory mechanism of subcellular mRNA localization. The maintenance of local transcriptomes contributes to the morphology and function of other cells with specialized subcellular regions, including *Drosophila* embryos, intestinal epithelia, and neurons (Gavis and Lehmann 1992; Moor et al. 2017; Perycz et al. 2011). These studies have discovered how RNA localization can be established through active trafficking, local anchoring, and selective stabilization, protection, or degradation. Many RNA-binding proteins are known; however, the interactions between RNA and RBPs depend on linear RNA sequences as well as the secondary structures formed by long mRNAs, making the computational prediction of RNA-RBP interactions unreliable. To this end, MPRAs are needed to provide reliable empirical data on the localization sequences that are necessary and sufficient in for RNA localization in a given cell type.

After our initial soma versus dendrite transcriptomic profiling by RNA-seq, we selected transcripts for further study based on their significant dendrite enrichment and for some, their predicted roles in cell projections and cytoskeletal organization. Of these, only *Samd4, Synpo*, and *Fgd3* have been studied in the skeleton. *Samd4* deficiency results in delayed bone formation and mineralization in mice by relieving repression of *Mig6* translation (Niu et al. 2017). *Synpo* was identified in a GWAS study for osteoporosis, and its knockdown reduced the osteogenic differentiation of human bone-derived mesenchymal stem cells (Lin and Pan 2021). A variant of *Fgd3* was identified as a risk trait in cattle with skeletal dysplasia (Takasuga et al. 2015), though it was primarily studied in growth plate cartilage. Ultimately, we aim to study the proper localization of these transcripts rather than the phenotype of their full ablation.

Our identification of precise localization sequences in osteocytes was limited to some degree by the inability to detect some 3’UTR oligos in the final MPRA analysis. This was most apparent in the 3’UTR of *Fgd3*, where 39 consecutive oligos did not meet the detection threshold for analysis in the MPRA, likely due to high GC content. The union of these oligos caused dendrite localization in followup studies, but we were unable to identify a peak sequence in this region with the same precision as the other 3’UTRs.

We were also unable to detect peaks of dendrite targeting activity within the 3’UTRs of *Tagln* and *Palld*. In the *Tagln* 3’UTR, seven oligos were not analyzed in the MPRA, but at most three were sequential, and the adjacent overlapping oligos were not significantly enriched in either compartment. In the *Palld* 3’UTR, six oligos were not analyzed in the MPRA, but their sequences were included in the larger fragments analyzed separately (Supp Figure 4). In these cases, we do not think that detection of oligos impacted our ability to identify localization sequences. We had anticipated that most localization information would be encoded in single, linear sequences in the range of 50-250 nt (Arora et al. 2022). However, prior work has also described 3’UTRs that broadly encode localization information in shorter sequences that additively leads to the full behavior of the 3’UTR (Mikl et al. 2022; Mendonsa et al. 2023). It is also possible that secondary structures like G-quadruplexes are formed from sequences that were never included in the same reporter constructs here (Goering et al. 2020). Further bioinformatic analysis or 3’UTR truncation experiments could be used to identify localization sequences for these transcripts. We also identified dendrite-enriched transcripts that were not regulated by their 3’UTRs, and additional work could investigate localization mechanisms driven by 5’UTRs, coding sequences, or stability.

We demonstrated localization of *Palld* mRNA and 18s rRNA in murine osteocytes *in situ*, suggesting that mRNA could be trafficked and locally translated *in vivo*. However, the majority of our current studies were conducted in a single osteocyte-like cell line, limiting our ability to assess the physiologic relevance of this mechanism in bone. Studies in neuronal cell lines, primary neurons, and micro-dissected rat brains have identified many commonalities across experimental systems; thus, we anticipate that proper regulation of mRNA localization will also have phenotypic consequences in osteocyte and bone biology.

In summary, here we describe that osteocytes possess a subcellular specific transcriptome within dendrites. We demonstrated local translation and identified necessary and sufficient localization sequences within the 3’UTRs. Our current findings provide a robust foundation to investigate novel RNA binding proteins (RBPs) that associate with newly-identified localization sequences and may control trafficking of a suite of dendrite enriched mRNAs. Furthermore, whether hormonal or mechanical signals that regulate osteocyte function impact dendrite mRNA trafficking and protein translation remains an exciting topic for future investigation.

## METHODS

### Cell culture and mechanical fractionation

Ocy454 osteocyte-like cells (Spatz et al. 2015; Wein et al. 2015) were cultured in MEM α supplemented with 10% FBS and 1% antibiotic/antimycotic (Gibco). Cells are grown in a 33ºC incubator with 5% CO2 until confluence, then moved to 37ºC to induce differentiation.

For fractionation experiments, we used a modified version of the protocol published by the Taliaferro group (Arora et al. 2021). Cells were grown in transwell membranes with 1 µm pores containing 2 mL media, each placed into one well of a deep six well companion plate containing 3 mL media (Corning #353102, #353502). Once confluent, cells were differentiated at 37ºC for 0-7 days.

To fractionate, media was aspirated and replaced by 1 mL PBS above the membrane and 2 mL PBS in the well. The top of the membrane was gently scraped to separate the soma fraction, which was collected into a conical tube on ice. The membrane was removed from the insert with a razor blade and placed into a clean six well plate containing 600 µL RNA or protein lysis buffer. A minimum of three membranes (half of a six-well plate) were combined for each replicate. Membranes incubated on a rocker at 4ºC for 15 minutes to digest the dendrite fraction. Soma suspensions in PBS were centrifuged at 4ºC and the pellets lysed in 600 µL RNA or protein lysis buffer for each soma fraction.

### Generation of reporter cell lines

To stably integrate reporter constructs, we genetically engineered Ocy454 cells to introduce an acceptor cassette into the *Rosa26* locus. We co-transfected an sgRNA targeting *Rosa26* (ACTCCAGTCTTTCTAGAAGA) and a plasmid expressing a repair template containing asymmetric LoxP sites (LoxP and Lox2272) flanking the blasticidin resistance gene (Khandelia, Yap, and Makeyev 2011). Following selection with blasticidin, we analyzed single cell clones to select a clonal cell line with a single insertion of the acceptor cassette. The resulting “LoxP osteocytes” were maintained in complete media with blasticidin at 5 µg/mL.

The reporter construct backbone used here is a derivative of the pRD-RIPE plasmid, which contains a doxycycline-inducible GFP reporter and puromycin resistance gene flanked by LoxP and Lox2272 sites (Khandelia, Yap, and Makeyev 2011). The Taliaferro group previously inserted a bi-directional doxycycline-inducible promoter driving Firefly luciferase and Renilla luciferase within the asymmetric LoxP sites (Arora et al. 2022; Goering et al. 2023). Reporter constructs can then be created by adding additional sequence at cut sites following the GFP coding sequence (BstXI) or Firefly luciferase coding sequence (PmeI), as described below.

LoxP osteocytes were plated at 250,000 cells/10 cm dish with blasticidin the day before transfection. To transfect, 5 µg reporter construct plasmid was mixed with 50 ng plasmid containing Cre recombinase in serum-free media with 15 uL PolyJet Reagent (SignaGen Laboratories). The mixture was added dropwise over one plate with fresh media without blasticidin. Media was changed the next day to remove the transfection reagents, and then again the next day to begin puromycin selection (4 µg/mL). Negative control transfections used no plasmid or Cre plasmid only. This procedure was repeated individually for each 3’UTR reporter plasmid to generate cell lines containing individual reporters. Cells with successful integration of the reporter plasmid were expanded and maintained in media containing puromycin.

### mRNA isolation and sequencing

For our first experiment, Ocy454 cells were plated onto transwell membranes and differentiated at 37ºC for 7 days. At fractionation, 5 wells of a 6-well plate were pooled into each of three replicates. Soma and dendrite lysate in RLT was passed through QIAshredder columns (Qiagen), then purified with PureLink RNA columns (Invitrogen) according to manufacturer’s instructions. Total RNA was submitted for library preparation and sequencing (BGI DNBseq platform).

At least 20 million paired-end reads per sample were obtained from PolyA-selected libraries, with >90% of reads mapping uniquely to the mouse transcriptome (GRCm39 Release 103; Salmon). Transcript counts were collapsed to gene counts, and differential expression analysis was performed with DESeq2 (Love, Huber, and Anders 2014). Genes with at least 10 normalized counts in all samples were considered for analysis.

### Generation of individual 3’UTR reporters

We selected the most abundant transcript of each dendrite-enriched mRNA chosen for followup. Ensembl 3’UTR sequences were used to order GBlocks (IDT) or to design primers to amplify the 3’UTR from Ocy454 dendrite cDNA. All 3’UTR inserts were flanked by overhangs allowing insertion following the FLuc coding sequence in the Taliaferro reporter plasmid. Purified plasmid backbone linearized with PmeI was assembled with 3’UTR inserts using InFusion Snap Assembly Master Mix (Takara #638947) according to manufacturer’s instructions. Products were transformed into chemically competent E. coli (NEB C2987H). Sanger sequencing confirmed accuracy of 3’UTR sequences in all final plasmid products.

### qRT-PCR and calculation of individual reporter localization

For localization experiments, LoxP osteocytes with stable integration of reporter constructs were grown in 6 well plates with porous membranes and differentiated at 37 °C for 7 days. To induce reporter plasmid expression, doxycycline (2 µg/mL) was added to culture media 48 hours before collection. Cells were then fractionated and mRNA isolated as described above, with three wells of a plate combined into each soma sample and paired dendrite sample. gDNA digestion and reverse transcription were performed with PrimeScript RT Reagent Kit (Takara RR047B). qPCR for endogenous transcripts, the coding sequences of FLuc (under control of the 3’UTR), and RLuc (internal control) were performed using SYBR green (Quantabio). To determine 3’UTR-driven reporter localization, FLuc expression was normalized to RLuc expression in each sample, then dendrites normalized to soma for each pair of samples. Finally, dendrite enrichment of FLuc for each 3’UTR was normalized to samples expressing no 3’UTR collected in each experiment.

### MPRA library design

Each 3’UTR sequence and the region immediately up and downstream was broken into 260 nucleotide (nt) fragments tiled at 4 nt intervals so that each nucleotide within the 3’UTR was represented in 65 unique oligonucleotides (oligos). Oligos were flanked by uniform PCR handles and synthesized by Twist Bioscience. The code for designing the MPRA oligos is available at https://github.com/TaliaferroLab/OligoPools/blob/master/makeoligopools/OligoPools_shortstep_260nt.py.

### MPRA library cloning

The oligonucleotide pool from Twist Bioscience was resuspended at 10 ng/µL. For each reporter (FLuc and GFP), we amplified 10 ng of oligo pool using Kapa HiFi HotStart DNA Polymerase (Kapa Biosystems, #KK2601) according to the manufacturer’s instructions. Amplification primers targeted the 20 nt PCR handles and added overhangs specific to the FLuc or GFP cloning site. PCR reactions were incubated with Exonuclease I for 2 hours at 37°C to digest single-stranded template, then purified using Zymo DNA Clean & Concentrator kits (#D4013).

The reporter plasmid backbone was linearized immediately following the FLuc or GFP reporter site using PmeI or BstXI, respectively, then gel extracted and purified. Oligo pools were cloned into the vector at a 6:1 molar ratio by Gibson Assembly at 50°C for 30 minutes (NEB E2611). The plasmid pools were ethanol precipitated, then 200 ng DNA were transformed into E. coli (DH10B Max chemically competent cells, Invitrogen). Transformed cells were spread across 25 15-cm agar-carbenicillin plates and incubated at 37 °C overnight. The next day, all colonies were scraped into LB media, and the plasmid library was purified using PureLink HiPure maxiprep columns (Invitrogen).

### MPRA

Osteocytes stably expressing the MPRA library were grown on microporous membranes and differentiated at 37°C for seven days, and doxycycline (2 µg/mL) was added to culture media 48 hours before collection. Cells were fractionated with six to nine wells of a plate combined into each soma and paired dendrite sample. gDNA digestion was performed on-column during RNA isolation (QIAgen 79254). 500 ng total RNA from each soma and dendrite sample was reverse transcribed for one hour at 55°C (SuperScript IV Reverse Transcriptase). Custom primers targeted the FLuc and GFP reporter sites and contained a partial Illumina read 1 primer sequence and an 8-nucleotide unique molecular identifier (UMI). Remaining RNA was then digested with 1 µl each of RNAse H and RNaseA/T1 mix for 30 minutes at 37°C.

Purified cDNA was amplified with Kapa HiFi HotStart DNA Polymerase (Kapa Biosystems, #KK2601) using 20 cycles for GFP and 25 cycles for FLuc reporter. Forward primers specific to the reporter site contained Illumina sequencing adaptors. Reverse primers targeted the partial Illumina read 1 sequence and contained the remaining sequencing adaptors and sample barcodes. PCR reactions were purified and size selected using double SPRI beads (Beckman, 0.55x, 0.8x). Libraries were sequenced by Novogene.

### MPRA analysis

More than 25 million paired-end reads per sample were trimmed using cutadapt to remove sequencing adapters, then aligned to the list of 3’UTR oligonucleotides using bowtie2. To minimize PCR bias, the number of unique UMIs (the first 8 nt of the reverse read) for each oligo was used as counts to calculate oligo abundance and dendrite enrichment (DESeq2). Oligos with at least 5 unique UMIs in all samples were used for statistical analysis (Supp Figure 3A). Custom code calculated the number of unique UMIs for each oligo: https://github.com/TaliaferroLab/OligoPools/blob/master/analyzeresults/UMIsperOligo.py.

### Puromycylation assays

To label nascent peptides, puromycin was added to cell culture media at 8 μg/mL at 37°C and then detected with anti-puromycin antibody (clone 12D10) (Schmidt et al. 2009). The translation elongation inhibitor cycloheximide (25 μM) was added 5 minutes before puromycin treatment as a control. For 2D images, Ocy454 cells were plated in chamber slides two days before puromycin treatment. For Western blots and 3D images, Ocy454 cells were grown to confluence on transwell membranes, then differentiated at 37°C for 0 or 2 days. For immunofluorescence, Ocy454 cells were fixed in 4% PFA, permeabilized in 0.05% saponin, then blocked in 1.5% BSA followed by rat anti-mouse Fc block (BD biosciences). Primary anti-puromycin antibody (clone 12D10, 10 μg/mL in 1.5% BSA/DPBS) was applied overnight, followed by secondary antibody and DAPI labeling. F-actin staining was performed during secondary antibody incubation (Abcam Cytopainter Green). Confocal imaging was performed on a Zeiss LSM 800 Airyscan confocal microscope with a 40x oil-immersion objective. Z-stacks were captured at 1 micron intervals and assembled into maximum intensity projections using ZEN microscopy software. For Western blots, whole cells or dendrites were digested from the membranes using TNT protein lysis buffer supplemented with NaF, vanadate, DTT, and protease inhibitor cocktail. Proteins were separated on 8% SDS-PAGE gels and labeled overnight with anti-puromycin antibody, then detected with HRP-linked anti-mouse secondary developed with ECL Plus (Thermo Scientific Pierce).

### Endoplasmic reticulum (ER) labeling

Ocy454 cells were transduced with lentivirus expressing mApple fused to ER protein calreticulin and KDEL ER-retention signal (gift from Dr. Ya-Cheng Liao). One week later, mApple-positive cells were collected by fluorescence activated cell sorting (BD FACSAria) and expanded. Cells were then grown in chamber slides for two days or on microporous membranes until day 7 of differentiation at 37 °C. Cells were fixed in 4% PFA, permeabilized in 0.05% saponin, and labeled for F-actin (Abcam Cytopainter Green).

### Deletion of endogenous RNA localization sequences

Two sgRNAs flanking each localization sequence were cloned into the px458 backbone (Addgene #48138, (Ran et al. 2013)) and transfected into Ocy454 cells with PolyJet Reagent (SignaGen Laboratories). Two days later, single GFP-expressing cells were isolated by fluorescence activated cell sorting (BD FACSAria) and expanded. Genomic DNA was isolated from clonal cell lines (QIAgen DNeasy Blood and Tissue Kit), and clones were screened for bi-allelic deletion by PCR.

### RNAscope

4 week old wild-type mice were euthanized with CO_2_ and hindlimbs quickly dissected. Tibias were fixed in 10% neutral buffered formalin at RT for 24 hours with gentle shaking. Decalcification was performed at 4 °C for 10 days in a solution of 9 parts 15% EDTA and 1 part 10% NBF, changed every 3 to 4 days. Bones were then washed in PBS, dehydrated in graded ethanols, and embedded in paraffin. Tissue sections were deparaffinized in xylene and two changes of 100% ethanol. Slides were then thoroughly dried in a 60°C oven and a hydrophobic barrier created around each tissue section. Sections were treated with hydrogen peroxide for 10 minutes at RT, pepsin for 30 minutes at 40 °C (Sigma R2283), and then hybridized with mRNA probes for 2 hours at 40 °C (ACD Bio). Amplification steps and DAB detection were performed according to kit instructions (RNAscope 2.5 HD Detection Reagent – Brown). Finally, sections were briefly counterstained with 50% hematoxylin, dehydrated, and mounted with coverslips.

### STARmap/RIBOmap

Spatially-resolved transcript amplicon readout mapping (STARmap) and ribosome-bound mRNA mapping (RIBOmap) were performed as previously described (X. Wang et al. 2018; Zeng et al. 2023). Briefly, Ocy454 cells cultured on type I collagen-coated, glass-bottom 24-well plates were fixed in 1.6% PFA and permeabilized with methanol. Next, cells were incubated overnight with a hybridization mixture containing probes specific to each mRNA of interest (IDT) (25 nM for Actb and Malat1, 50 nM for osteocyte gene targets). For STARmap, signal is generated upon hybridization of both primer and barcoded padlock probes. For RIBOmap, a splint probe targeting 18S rRNA is required to form the tri-component hybridization complex, thereby detecting only ribosome-bound mRNA. After washing, hybridized padlock probes were circularized for two hours and then amplified through rolling circle amplification for two hours. For detection, cells were incubated with fluorescently-conjugated oligos complementary to the amplified barcode region overnight at room temperature. Finally, nuclei were labeled with DAPI and cells were imaged in PBST on a Leica confocal microscope with a 40x oil immersion objective.

## Acknowledgements

We thank Lauren Surface, all members of the MGH Endocrine Unit, and the Wein laboratory for helpful discussions. We also thank Charlie Moffatt for expert technical input. M.N.W. acknowledges funding support from the National Institute of Health (R01DK116716), the Smith Family Foundation Odyssey Award, and the Chen Institute Massachusetts General Hospital Research Scholar (2024-2029) award. C.M.M. acknowledges support from the NIH (T32DK007028, F32AR081660). J.M.T. acknowledges support from the NIH (R35-GM133385). M.N.W. and C.M.M. acknowledge generous support from Louise Pearl Corman, Ph.D. Bone histology was performed by with Center for Skeletal Research at Massachusetts General Hospital, a NIH-funded program (P30 AR066261 and AR075042). Confocal microscopy was supported by the NIH Shared Instrumentation Grant (SIG) S10OD021577.

## Conflict of Interests

The authors declare competing interest. M.N.W. has received research funding from Radius Health for work unrelated to this project, and is a co-inventor on pending patents regarding the use of SIK inhibitors for osteoporosis. MNW serves as a consultant for Relation Therapeutics, Sitryx Therapeutics, Aditum Bio, and GLG.

## FIGURE LEGENDS

**Supplemental Figure 1:**
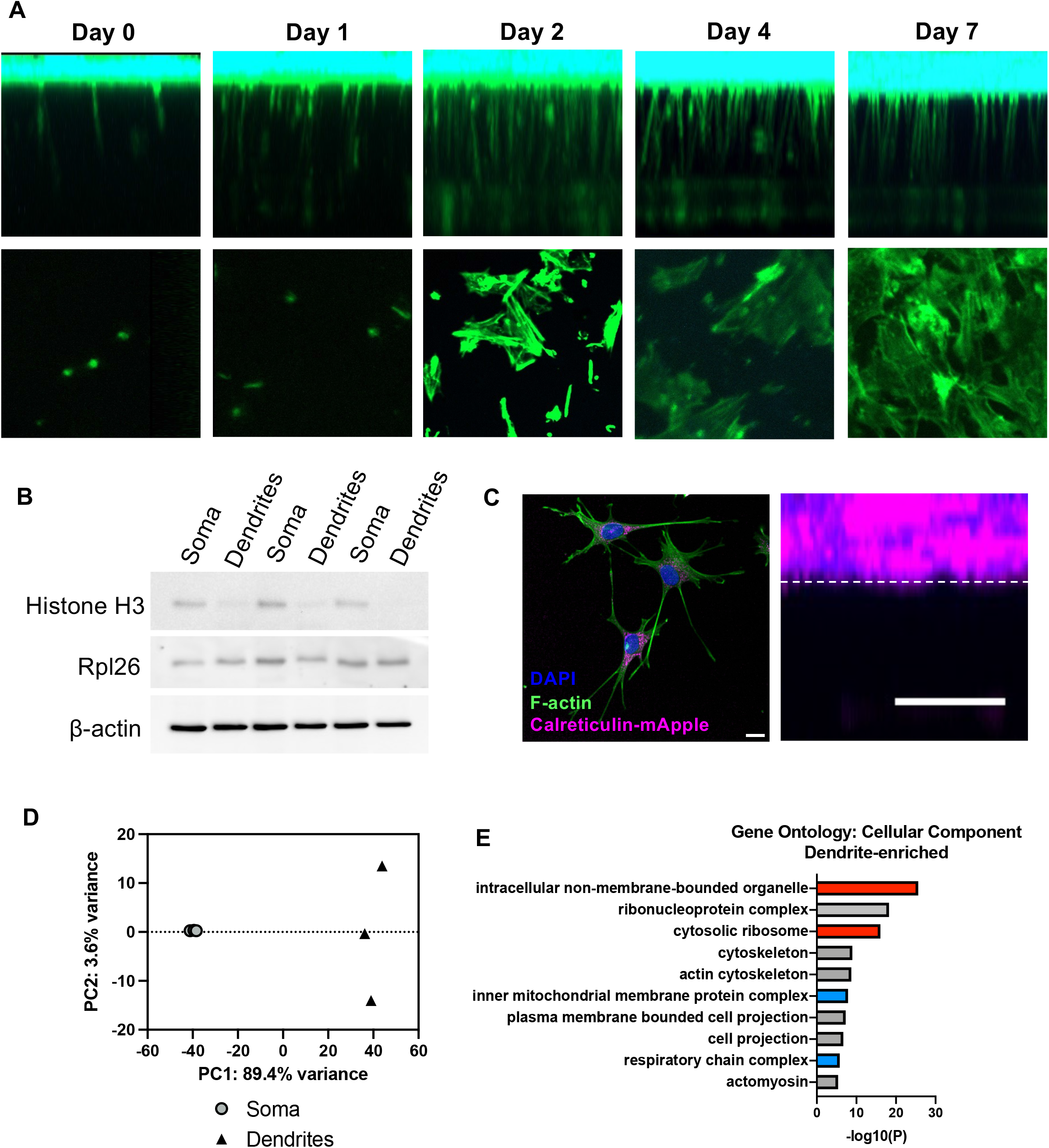
A) Side view of Z-stack images of dendrites extending through microporous membranes (top) and bottom view of membranes as dendrites emerge from pores (bottom) at the indicated differentiation timepoints. B) Immunoblots from paired soma and dendrite fractions. C) Ocy454 cells expressing mApple fused to ER protein calreticulin and KDEL ER-retention signal (magenta pseudocolored) were grown in 2D and labeled with phalloidin and DAPI (left, scale bar = 20 μm) or grown on microporous membranes (right, scale bar = 10 μm). D) Principal component analysis of transcript abundance in fractionated dendrites/soma. E) Gene ontology enrichments for 420 dendrite-enriched genes relative to 11,880 genes detected in all samples.

**Supplemental Figure 2:**
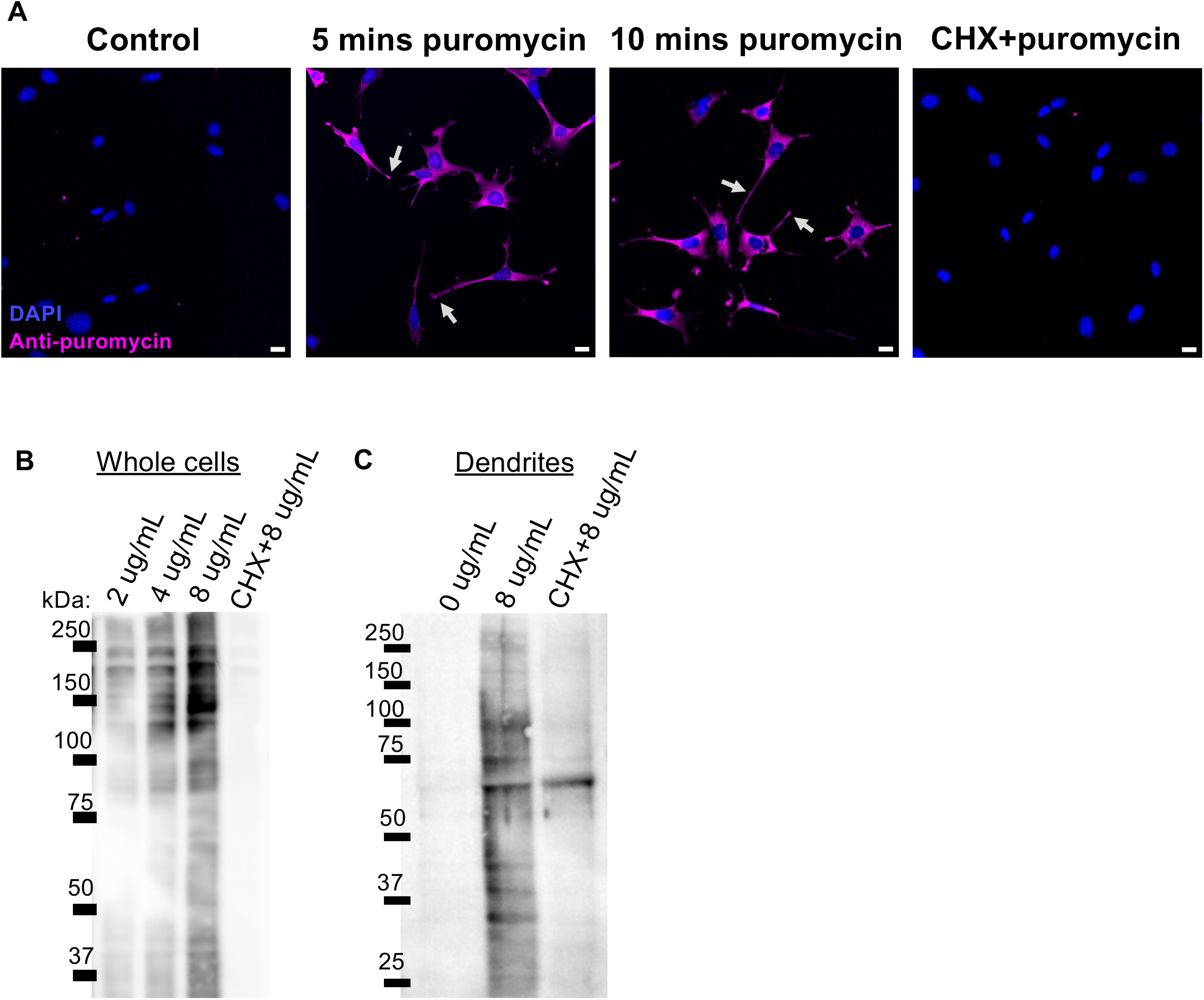
A) Anti-puromycin immunofluorescence (magenta) after indicated treatments in Ocy454 cells grown in 2D. Arrows indicate puromycylated peptides in dendrites. Cycloheximide (CHX) pre-treatment for 5 minutes blocks translational elongation and incorporation of puromycin into nascent peptides. Scale bars = 20 μm. B) Immunoblots in whole cells show increasing puromycylated proteins with increasing doses of puromycin after 10 minutes of treatment. Cycloheximide (CHX) pre-treatment for 5 minutes blocks translational elongation and incorporation of puromycin into nascent peptides. C) Immunoblots from dendrites at day 0 of differentiation that were treated with indicated doses of puromycin for 10 minutes after removal of soma.

**Supplemental Figure 3:**
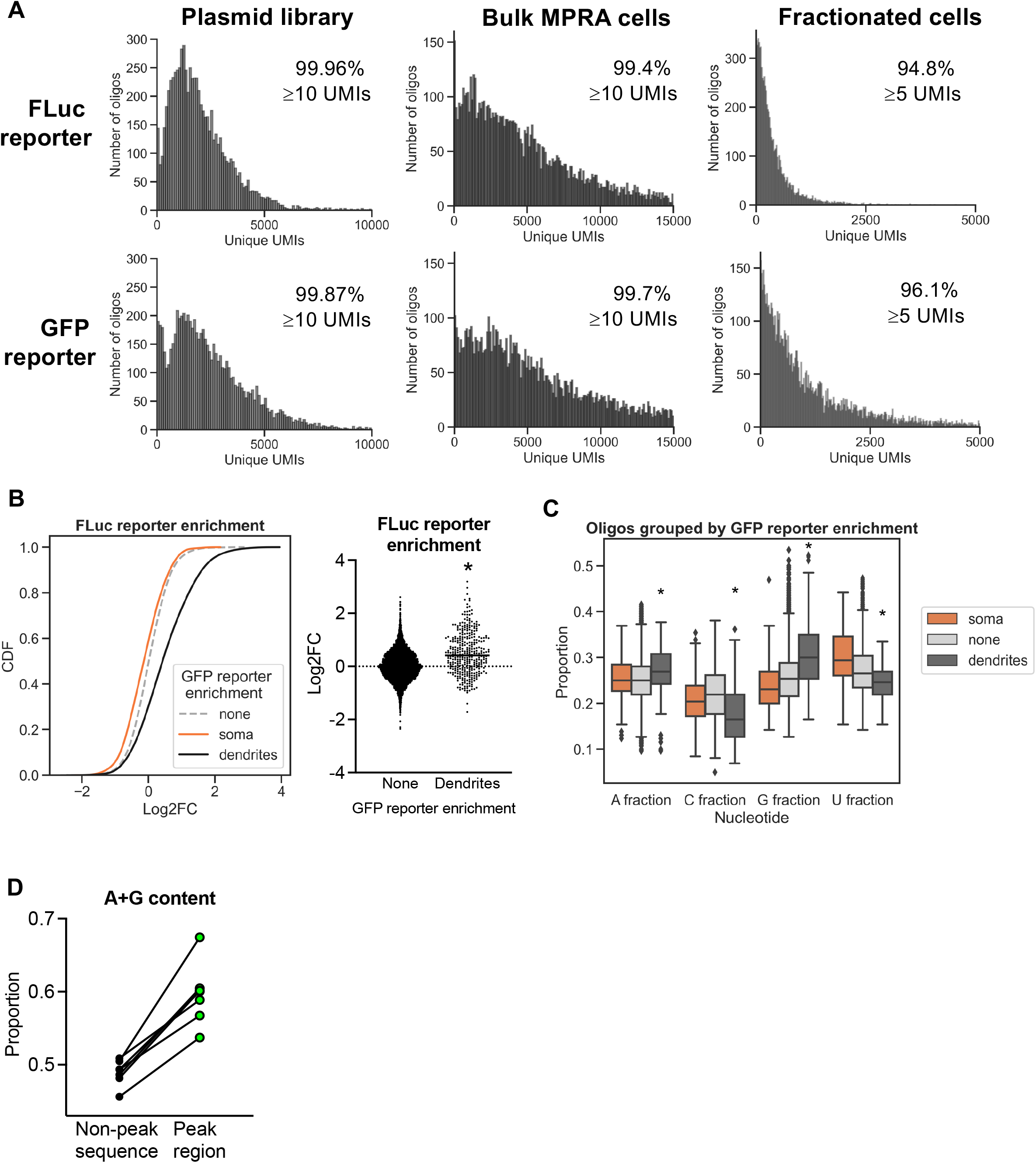
A) Distribution of oligonucleotide abundances at the FLuc and GFP reporters in the final constructed plasmid library, integrated into Ocy454 cells, and detected in the final fractionated samples (presented as mean of 3 soma and 3 dendrite samples). B) Concordance of results across reporter contexts. Dendrite enrichment of oligos at the FLuc reporter site as a function of their enrichment at the GFP reporter site. *p<0.0001. C) Nucleotide content of oligos as a function of their enrichment at the GFP reporter site. *p<0.0001 dendrite-enriched vs no enrichment. D) The proportion of A+G nucleotides in the peak region compared to non-peak sequence from each 3’UTR. Descending order of their peak region A+G content: *Cgnl1, Synpo, Net1, Trak2, Samd4, Sh3pxd2a, Acta2*.

**Supplemental Figure 4:**
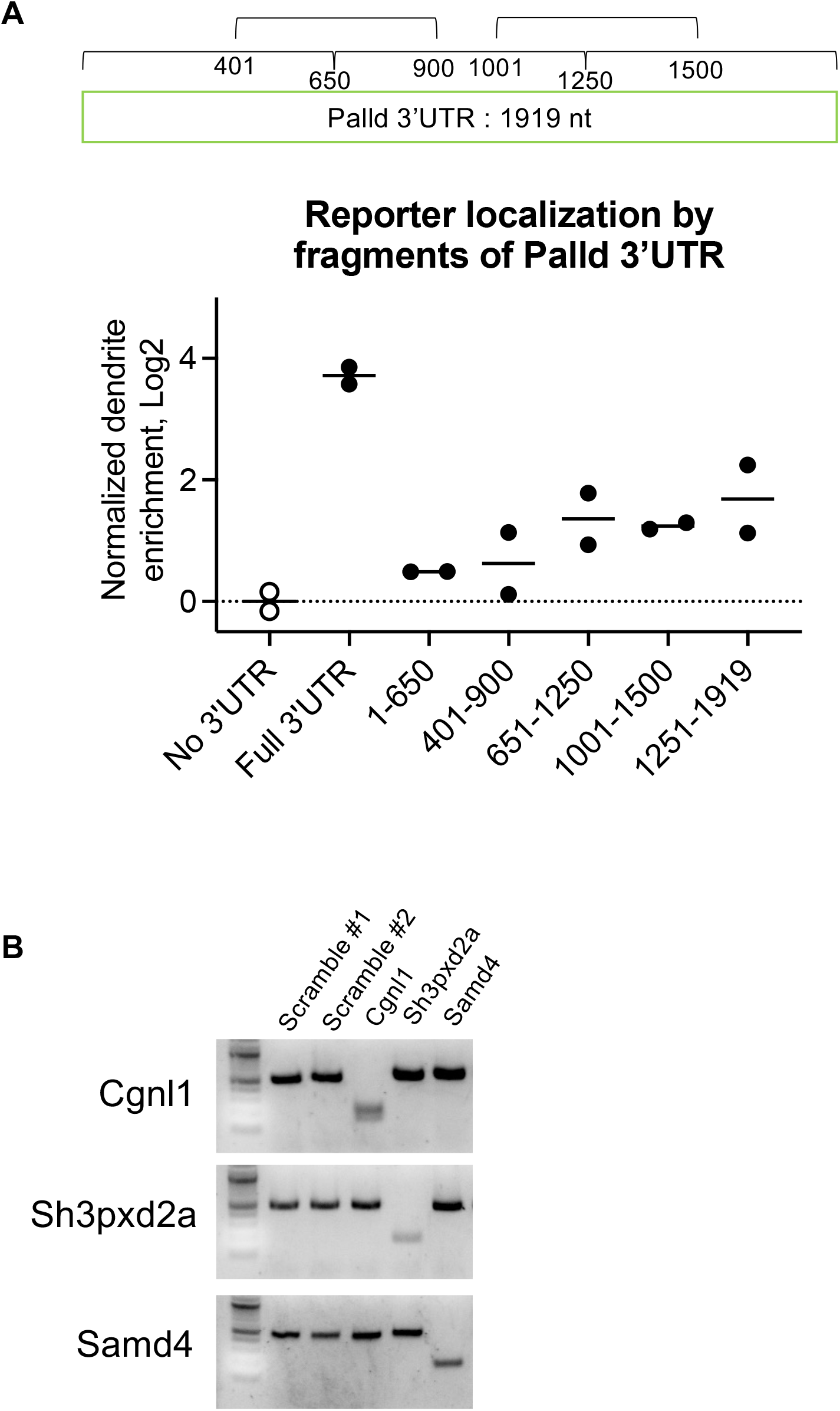
A) Dendrite enrichment of FLuc fused to fragments of the *Palld* 3’UTR spanning the indicated nucleotides relative to RLuc. The localization of FLuc without any added localization sequence was used as a normalization control. B) PCR products indicate specific deletion of approximately 300 bp from the 3’UTRs of Cgnl1, Sh3pxd2a, and Samd4 in individual clonal cell lines. Scramble cell lines received two non-targeting sgRNAs.

## REFERENCES

Arora, Ankita, Roberto Castro-Gutierrez, Charlie Moffatt, Davide Eletto, Raquel Becker, Maya Brown, Andreas E. Moor, Holger A. Russ, and J. Matthew Taliaferro. 2022. “High-Throughput Identification of RNA Localization Elements in Neuronal Cells.” Nucleic Acids Research 50 (18): 10626–42. 10.1093/nar/gkac763.

Arora, Ankita, Raeann Goering, Hei-Yong G. Lo, and Matthew J. Taliaferro. 2021. “Mechanical Fractionation of Cultured Neuronal Cells into Cell Body and Neurite Fractions.” Bio-Protocol 11 (11): e4048. 10.21769/BioProtoc.4048.

Biever, Anne, Caspar Glock, Georgi Tushev, Elena Ciirdaeva, Tamas Dalmay, Julian D. Langer, and Erin M. Schuman. 2020. “Monosomes Actively Translate Synaptic mRNAs in Neuronal Processes.” Science (New York, N.Y.) 367 (6477): eaay4991. 10.1126/science.aay4991.

Chekulaeva, Marina. 2024. “Mechanistic Insights into the Basis of Widespread RNA Localization.” Nature Cell Biology 26 (7): 1037–46. 10.1038/s41556-024-01444-5.

Dallas, Sarah L., Matthew Prideaux, and Lynda F. Bonewald. 2013. “The Osteocyte: An Endocrine Cell … and More.” Endocrine Reviews 34 (5): 658–90. 10.1210/er.2012-1026.

Dole, Neha S., Courtney M. Mazur, Claire Acevedo, Justin P. Lopez, David A. Monteiro, Tristan W. Fowler, Bernd Gludovatz, et al. 2017. “Osteocyte-Intrinsic TGF-β Signaling Regulates Bone Quality through Perilacunar/Canalicular Remodeling.” Cell Reports 21 (9): 2585–96. 10.1016/j.celrep.2017.10.115.

Engel, Krysta L., Ankita Arora, Raeann Goering, Hei-Yong G. Lo, and J. Matthew Taliaferro. 2020. “Mechanisms and Consequences of Subcellular RNA Localization across Diverse Cell Types.” Traffic (Copenhagen, Denmark) 21 (6): 404–18. 10.1111/tra.12730.

Fowler, Tristan W., Claire Acevedo, Courtney M. Mazur, Faith Hall-Glenn, Aaron J. Fields, Hrishikesh A. Bale, Robert O. Ritchie, Jeffrey C. Lotz, Thomas P. Vail, and Tamara Alliston. 2017. “Glucocorticoid Suppression of Osteocyte Perilacunar Remodeling Is Associated with Subchondral Bone Degeneration in Osteonecrosis.” Scientific Reports 7 (March):44618. 10.1038/srep44618.

Gavis, E. R., and R. Lehmann. 1992. “Localization of Nanos RNA Controls Embryonic Polarity.” Cell 71 (2): 301–13. 10.1016/0092-8674(92)90358-j.

Goering, Raeann, Ankita Arora, Megan C. Pockalny, and J. Matthew Taliaferro. 2023. “RNA Localization Mechanisms Transcend Cell Morphology.” eLife 12 (March):e80040. 10.7554/eLife.80040.

Goering, Raeann, Laura I. Hudish, Bryan B. Guzman, Nisha Raj, Gary J. Bassell, Holger A. Russ, Daniel Dominguez, and J. Matthew Taliaferro. 2020. “FMRP Promotes RNA Localization to Neuronal Projections through Interactions between Its RGG Domain and G-Quadruplex RNA Sequences.” eLife 9 (June):e52621. 10.7554/eLife.52621.

Khandelia, Piyush, Karen Yap, and Eugene V. Makeyev. 2011. “Streamlined Platform for Short Hairpin RNA Interference and Transgenesis in Cultured Mammalian Cells.” Proceedings of the National Academy of Sciences of the United States of America 108 (31): 12799–804. 10.1073/pnas.1103532108.

Lin, Bingyuan, and Zhijun Pan. 2021. “Consensus Gene Modules Related to Levels of Bone Mineral Density (BMD) among Smokers and Nonsmokers.” Bioengineered 12 (2): 10134–46. 10.1080/21655979.2021.2000746.

Love, Michael I., Wolfgang Huber, and Simon Anders. 2014. “Moderated Estimation of Fold Change and Dispersion for RNA-Seq Data with DESeq2.” Genome Biology 15 (12): 550. 10.1186/s13059-014-0550-8.

Mazur, Courtney M., Jonathon J. Woo, Cristal S. Yee, Aaron J. Fields, Claire Acevedo, Karsyn N. Bailey, Serra Kaya, et al. 2019. “Osteocyte Dysfunction Promotes Osteoarthritis through MMP13-Dependent Suppression of Subchondral Bone Homeostasis.” Bone Research 7:34. 10.1038/s41413-019-0070-y.

Mendonsa, Samantha, Nicolai von Kügelgen, Sayaka Dantsuji, Maya Ron, Laura Breimann, Artem Baranovskii, Inga Lödige, et al. 2023. “Massively Parallel Identification of mRNA Localization Elements in Primary Cortical Neurons.” Nature Neuroscience 26 (3): 394–405. 10.1038/s41593-022-01243-x.

Middleton, Sarah A., James Eberwine, and Junhyong Kim. 2019. “Comprehensive Catalog of Dendritically Localized mRNA Isoforms from Sub-Cellular Sequencing of Single Mouse Neurons.” BMC Biology 17 (1): 5. 10.1186/s12915-019-0630-z.

Mikl, Martin, Davide Eletto, Malak Nijim, Minkyoung Lee, Atefeh Lafzi, Farah Mhamedi, Orit David, Simona Baghai Sain, Kristina Handler, and Andreas E. Moor. 2022. “A Massively Parallel Reporter Assay Reveals Focused and Broadly Encoded RNA Localization Signals in Neurons.” Nucleic Acids Research 50 (18): 10643–64. 10.1093/nar/gkac806.

Moharrer, Yasaman, and Joel D. Boerckel. 2021. “Tunnels in the Rock: Dynamics of Osteocyte Morphogenesis.” Bone 153 (December):116104. 10.1016/j.bone.2021.116104.

Moor, Andreas E., Matan Golan, Efi E. Massasa, Doron Lemze, Tomer Weizman, Rom Shenhav, Shaked Baydatch, et al. 2017. “Global mRNA Polarization Regulates Translation Efficiency in the Intestinal Epithelium.” Science (New York, N.Y.) 357 (6357): 1299–1303. 10.1126/science.aan2399.

Niu, Ningning, Jian-Feng Xiang, Qin Yang, Lijun Wang, Zhanying Wei, Ling-Ling Chen, Li Yang, and Weiguo Zou. 2017. “RNA-Binding Protein SAMD4 Regulates Skeleton Development through Translational Inhibition of Mig6 Expression.” Cell Discovery 3:16050. 10.1038/celldisc.2016.50.

Perycz, Malgorzata, Anna S. Urbanska, Pawel S. Krawczyk, Kamil Parobczak, and Jacek Jaworski. 2011. “Zipcode Binding Protein 1 Regulates the Development of Dendritic Arbors in Hippocampal Neurons.” The Journal of Neuroscience: The Official Journal of the Society for Neuroscience 31 (14): 5271–85. 10.1523/JNEUROSCI.2387-10.2011.

Ran, F. Ann, Patrick D. Hsu, Jason Wright, Vineeta Agarwala, David A. Scott, and Feng Zhang. 2013. “Genome Engineering Using the CRISPR-Cas9 System.” Nature Protocols 8 (11): 2281–2308. 10.1038/nprot.2013.143.

Schmidt, Enrico K., Giovanna Clavarino, Maurizio Ceppi, and Philippe Pierre. 2009. “SUnSET, a Nonradioactive Method to Monitor Protein Synthesis.” Nature Methods 6 (4): 275–77. 10.1038/nmeth.1314.

Spatz, Jordan M., Marc N. Wein, Jonathan H. Gooi, Yili Qu, Jenna L. Garr, Shawn Liu, Kevin J. Barry, et al. 2015. “The Wnt Inhibitor Sclerostin Is Up-Regulated by Mechanical Unloading in Osteocytes in Vitro.” The Journal of Biological Chemistry 290 (27): 16744–58. 10.1074/jbc.M114.628313.

Staines, Katherine A., Behzad Javaheri, Peter Hohenstein, Robert Fleming, Ekele Ikpegbu, Erin Unger, Mark Hopkinson, David J. Buttle, Andrew A. Pitsillides, and Colin Farquharson. 2017. “Hypomorphic Conditional Deletion of E11/Podoplanin Reveals a Role in Osteocyte Dendrite Elongation.” Journal of Cellular Physiology 232 (11): 3006–19. 10.1002/jcp.25999.

Takasuga, Akiko, Kunio Sato, Ryouichi Nakamura, Yosuke Saito, Shinji Sasaki, Takehito Tsuji, Akio Suzuki, et al. 2015. “Non-Synonymous FGD3 Variant as Positional Candidate for Disproportional Tall Stature Accounting for a Carcass Weight QTL (CW-3) and Skeletal Dysplasia in Japanese Black Cattle.” PLoS Genetics 11 (8): e1005433. 10.1371/journal.pgen.1005433.

Taliaferro, J. Matthew, Marina Vidaki, Ruan Oliveira, Sara Olson, Lijun Zhan, Tanvi Saxena, Eric T. Wang, et al. 2016. “Distal Alternative Last Exons Localize mRNAs to Neural Projections.” Molecular Cell 61 (6): 821–33. 10.1016/j.molcel.2016.01.020.

Tiede-Lewis, LeAnn M., Yixia Xie, Molly A. Hulbert, Richard Campos, Mark R. Dallas, Vladimir Dusevich, Lynda F. Bonewald, and Sarah L. Dallas. 2017. “Degeneration of the Osteocyte Network in the C57BL/6 Mouse Model of Aging.” Aging 9 (10): 2190–2208. 10.18632/aging.101308.

Torre, E. R., and O. Steward. 1992. “Demonstration of Local Protein Synthesis within Dendrites Using a New Cell Culture System That Permits the Isolation of Living Axons and Dendrites from Their Cell Bodies.” The Journal of Neuroscience: The Official Journal of the Society for Neuroscience 12 (3): 762–72. 10.1523/JNEUROSCI.12-03-00762.1992.

Wang, Jialiang S., Tushar Kamath, Courtney M. Mazur, Fatemeh Mirzamohammadi, Daniel Rotter, Hironori Hojo, Christian D. Castro, et al. 2021. “Control of Osteocyte Dendrite Formation by Sp7 and Its Target Gene Osteocrin.” Nature Communications 12 (1): 6271. 10.1038/s41467-021-26571-7.

Wang, Jialiang S., and Marc N. Wein. 2022. “Pathways Controlling Formation and Maintenance of the Osteocyte Dendrite Network.” Current Osteoporosis Reports 20 (6): 493–504. 10.1007/s11914-022-00753-8.

Wang, Xiao, William E. Allen, Matthew A. Wright, Emily L. Sylwestrak, Nikolay Samusik, Sam Vesuna, Kathryn Evans, et al. 2018. “Three-Dimensional Intact-Tissue Sequencing of Single-Cell Transcriptional States.” Science (New York, N.Y.) 361 (6400): eaat5691. 10.1126/science.aat5691.

Wein, Marc N., Jordan Spatz, Shigeki Nishimori, John Doench, David Root, Philip Babij, Kenichi Nagano, et al. 2015. “HDAC5 Controls MEF2C-Driven Sclerostin Expression in Osteocytes.” Journal of Bone and Mineral Research: The Official Journal of the American Society for Bone and Mineral Research 30 (3): 400–411. 10.1002/jbmr.2381.

Zappulo, Alessandra, David van den Bruck, Camilla Ciolli Mattioli, Vedran Franke, Koshi Imami, Erik McShane, Mireia Moreno-Estelles, et al. 2017. “RNA Localization Is a Key Determinant of Neurite-Enriched Proteome.” Nature Communications 8 (1): 583. 10.1038/s41467-017-00690-6.

Zeng, Hu, Jiahao Huang, Jingyi Ren, Connie Kangni Wang, Zefang Tang, Haowen Zhou, Yiming Zhou, et al. 2023. “Spatially Resolved Single-Cell Translatomics at Molecular Resolution.” Science (New York, N.Y.) 380 (6652): eadd3067. 10.1126/science.add3067.

